# Structural, genetic, and adaptive basis of superhydrophobicity in rice leaves

**DOI:** 10.1101/2024.12.05.626993

**Authors:** Asuka Hiraiwa, Saori Aiga, Tian Zhu, Manaki Mimura, Takanori Yoshikawa, Yutaka Sato, Jun-ichi Itoh

## Abstract

In most plants, leaf surfaces exhibit water-repellent properties, which protect against pathogens and weather. Plants such as lotus and rice have evolved outstanding water-repellent properties, so-called superhydrophobicity. However, the structural and genetic basis of these properties in crop plants remains unclear. In this study, we identified the genes and microstructures responsible for superhydrophobicity by analyzing water repellence in rice mutants and cultivars, as well as wild *Oryza* species. Analysis of the surface structure of mutant leaves revealed that the degree of water repellency in rice largely depends on the accumulation of cuticular wax. Meanwhile, although papillae differentiation contributes minorly to water repellency, it is suggested to play a crucial role in achieving superhydrophobicity. These morphological characteristics are regulated by at least five previously characterized genes. We also found that superhydrophobicity is present in ancestral species, is retained by most cultivated rice, and appears to have evolved independently of rice domestication. Understanding the mechanisms of water repellence and their applications should provide important insight into the natural design of water repellence, as well as a basis for future innovation.

## Introduction

In plants, the surface cuticle is responsible for plant–environment interactions and plays an important role in defense against environmental and biological stresses such as drought, ultraviolet radiation, pathogens, and insects (Koch and Ensikat, 2008;Yeats and Rose, 2013; Barthlott et al., 2017; Jolliffe et al., 2023). One of the physical properties associated with the cuticle is water repellence. The degree of water repellence varies among plant species, but most plants exhibit very good water-repellent properties (Barthlott et al., 2017). Water repellence facilitates gas exchange through stomata by maintaining contact with the atmosphere when rainwater adheres to leaf surfaces. It also hinders the attachment of pathogens to the epidermis, thereby conferring protection against disease (Barthlott et al., 2017).

Epicuticular wax deposited on the leaf surface is an important contributor to water repellence (Koch and Ensikat, 2008; Yeats and Rose, 2013; Barthlott et al., 2017; Jolliffe et al., 2023). This epicuticular wax consists of compounds derived from very-long-chain fatty acids (VLCFAs; C20–C34), including alkanes, aldehydes, primary and secondary alcohols, ketones, and esters, although the precise composition varies for different plant species, organs, growth stages, and conditions (Yeats and Rose, 2013; Lee and Suh, 2015; Batsale et al., 2021). VLCFAs are synthesized in the endoplasmic reticulum by fatty acid elongation (FAE) complex enzymes, including β-ketoacyl-CoA synthase (KCS), β-ketoacyl-CoA reductase (KCR), β-hydroxy acyl-CoA dehydratase, and enoyl-CoA reductase (Kunst and Samuels, 2009; Lee and Suh, 2015). The elongated VLCFA-CoA is processed by an alkane-forming or alcohol-forming pathway to generate a cuticular wax mixture, which is eventually secreted (Kunst and Samuels, 2009); (Yeats and Rose, 2013; Lee and Suh, 2015). The secreted cuticular wax forms epicuticular crystals that are deposited on the surface of plants (Koch and Ensikat, 2008). These wax crystals may be crucial for leaf-surface water repellence (Bhushan, 2012; Barthlott et al., 2017).

Some plants exhibit outstanding water-repellent properties associated with additional functions, so-called superhydrophobicity. Lotus (*Nelumbo nucifera*) is one such species (Guo and Liu, 2007; Bhushan, 2012; Barthlott et al., 2017). Superhydrophobic leaf surfaces have micro-protrusions and wax crystals (Barthlott and Neinhuis, 1997; Barthlott et al., 2017) that create static contact angles on the leaf surface greater than 150°. In a plant species with a superhydrophobic surface combined with low hysteresis, droplets of water roll across the surface and wash away dirt. This self-cleaning property maintains gas exchange through stomata, increases photosynthetic efficiency, and is called “the lotus effect” (Barthlott and Neinhuis, 1997; Neinhuis and Barthlott, 1997; Bhushan, 2012; Barthlott et al., 2017; Yeats and Rose, 2013). Rice plants also exhibit superhydrophobicity, wax crystals on the leaves, and differentiated papillae, which are micro-protrusions on the epidermis (Neinhuis and Barthlott, 1997; Guo and Liu, 2007; Barthlott et al., 2017). Rice is often grown in paddy fields, and water repellence is important because it generates a gas film across the submerged epidermis (Raven, 2008; Kurokawa et al., 2018).

The genes necessary for water repellence may be identified by analyzing mutants that are deficient in this property. One requirement for water repellence may be the synthesis of cuticular wax. Mutants deficient in the accumulation of such wax have been reported in various plant species (Yeats and Rose, 2013; Lee and Suh, 2015). In rice, several mutants with abnormal accumulation of cuticular wax exhibit reduced tolerance to environmental stress, as well as reduced water repellence (Yu et al., 2008; Islam et al., 2009; Qin et al., 2011; Mao et al., 2012; Gan et al., 2016; Zhang et al., 2016; Gan et al., 2017; Wang et al., 2017; Kurokawa et al., 2018). Among these, *crystal-sparse leaf* (*wsl*) mutants show reduced accumulation of epicuticular wax and abnormal cuticular wax composition. *WAX CRYSTAL-SPARSE LEAF1* (*WSL1*) encodes a KCS that catalyzes the biosynthesis of C24 from C20 VLCFAs (Yu et al., 2008). Another KCS, encoded by *WSL4*, is involved in the elongation of long (>C22) VLCFAs (Gan et al., 2017; Wang et al., 2017). *WSL3* encodes a KCR for a FAE complex that is essential for the elongation of VLCFAs and accumulation of leaf wax (Gan et al., 2016). *OsGL1–1/WSL2* is a homolog of the *Arabidopsis WAX2* and maize *GL1* gene, which does not encode a component of an FAE complex. *OsGL1–1/WSL2* affects epicuticular wax production by participating in the elongation of VLCFAs, and mutants show reduced water repellence (Qin et al., 2011; Mao et al., 2012). *OsHSD1*, a member of the hydroxysteroid dehydrogenases gene family in rice, is also involved in leaf epicuticular wax production and lipid homeostasis (Zhang et al., 2016). *Leaf Gas Film 1* (*LGF1*), the same gene as *OsHSD1*, is necessary for facilitating plant gas exchange under water (Kurokawa et al., 2018). The *dripping wet leaf 7* (*drp7*) mutant of *LGF1* cannot maintain a gas film and exhibits reduced water repellence. *LGF1* regulates C30 primary alcohol synthesis, which is necessary to produce an abundance of epicuticular wax on leaf surfaces (Kurokawa et al., 2018).

Another factor that may be important for water repellence is the microstructure of leaf epidermal cells. Little is known about the underlying genetic mechanisms that create the microstructure of epidermal cells, but *BRIGHT GREEN LEAF* (*BGL*) is involved in the formation of papillae on rice leaves. *BGL* encodes OsRopGEF10, a member of the plant-specific RopGEF family of proteins, but its influence on water repellence has not been examined (Yoo et al., 2011).

Water repellence in plants is influenced by the physical properties of the leaf epidermis but the factors involved may be very complex (Koch and Ensikat, 2008; Bhushan, 2012; Barthlott et al., 2017). Little is known about the determinants and genes that are important for water repellence during plant growth. In this study, we analyzed the relationship between leaf surface structure and water repellence in rice. We used various mutants, cultivars, and wild *Oryza* species to understand how superhydrophobicity is achieved. Our results provide valuable insight that may be used to influence water repellence in plants.

## Results

### Structure of different parts of the leaf surface and water repellence

To understand the relationship between leaf surface structure and water repellence, we studied four different parts of the rice (*O*. *sativa*) leaf using SEM. On the adaxial surface of the leaf blade, there were many uniformly distributed 1–2 µm papillary protrusions on the epidermal cell layer (Fig. 1A). The leaf surface was also covered with wax crystals (inset). On the abaxial surface of the leaf blade, in addition to papillae that were similar in size to those on the adaxial surface, papillae larger than 20 µm were also observed (Fig. 1B). Neither papillae nor wax crystals were observed on the adaxial surface of the leaf sheath (Fig. 1C), whereas on the abaxial surface of the leaf sheath, there were small papillae (approximately 1.5 µm; Fig. 1D).

**Figure 1.**
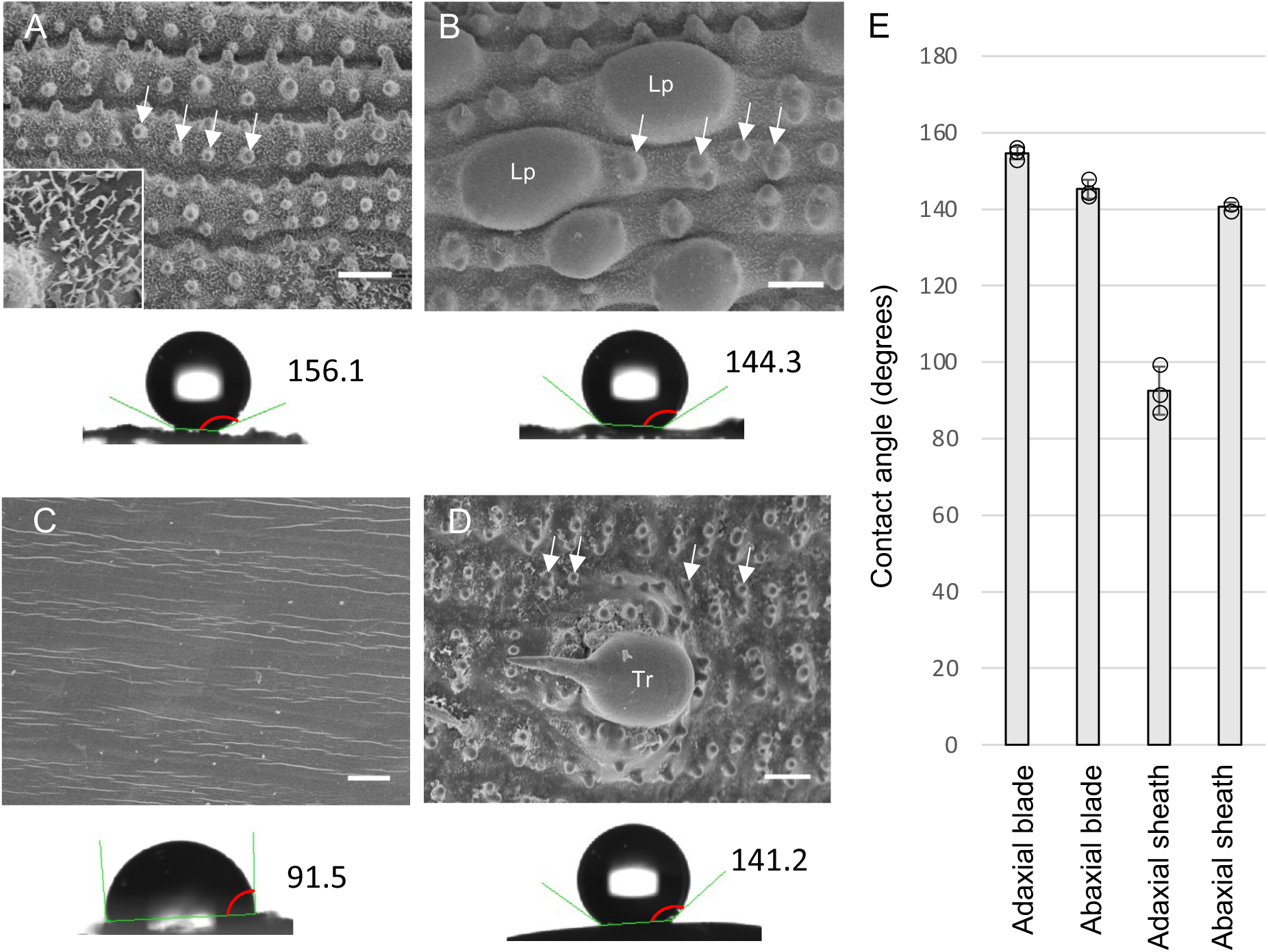
Surface structure and water repellence for different parts of the leaf. **A)** Scanning electron microscopy image of the adaxial surface of the 6th leaf blade of a rice plant. Inset: an enlarged view of the surface. **B)** Abaxial surface of the 6th leaf blade. **C)** Adaxial surface of the 6th leaf sheath. **D)** Abaxial surface of the 6th leaf sheath. For each panel, the measurement of the contact angle of the representative sample is indicated. The red arc indicates the contact angle. White arrows in **A**, **B**, and **D** indicate epidermal papillae. Lp: large papilla; Tr: trichome. Scale bars in (**A**–**D**) = 10 µm. **F)** Contact angles of the leaf parts in rice. Error bars indicate standard deviation (*n* = 3).

A contact angle meter was used to quantify water repellence on these leaf surfaces (Fig. 1A–D bottom). Greater contact angles indicate superior water repellence. In particular, a surface exhibiting contact angles > 150° is considered superhydrophobic (Barthlott et al., 2017; Koch and Barthlott, 2009). Under different experimental conditions, the contact angles for rice leaves are reportedly 157 ± 2° and 162° (Barthlott and Neinhuis, 1997; Neinhuis and Barthlott, 1997). Under our experimental conditions, the contact angle for the adaxial leaf blade was 155°, which represents the greatest value among the four parts of the 6th leaf; the smallest value was for the adaxial surface of the leaf sheath (93°). The values for the abaxial surface of the leaf blade and the leaf sheath were 145° and 141°, respectively (Fig. 1E). These results indicate that the degree of water repellence differs on different parts of the rice leaf, and the adaxial surface of the leaf blade, which has uniformly distributed papillae and wax crystals, exhibits the greatest water repellence and superhydrophobicity.

### Changes in leaf surface structure during growth and development

To investigate how these microstructures are formed, the surface of the 6th leaf primordia of wild-type plastochron4 (P4) was examined 20 days after germination. Various parts of the leaf from the base to the apical axis were observed in plastic sections using SEM. The surface of the immature epidermal cell layer at the base of the leaf blade had no papillae or wax crystals (Fig. 2A, D). On the epidermal cells at the center of the leaf blade, there were multiple small protrusions (papillae primordia) but no wax crystals (Fig. 2B, E). On the surface of the apical part of the leaf blade, almost complete papillae and wax crystals were present. In addition, when toluidine blue was applied, the epidermal cells were clearly stained, suggesting a thickening of the cuticle here (Fig. 2C, F). These results indicate that the leaf microstructure that is responsible for water repellence develops gradually from the apical part of the leaf blade toward the base in the P4 leaf primordium. Papillae form first, followed by the accumulation of epicuticular wax.

**Figure 2.**
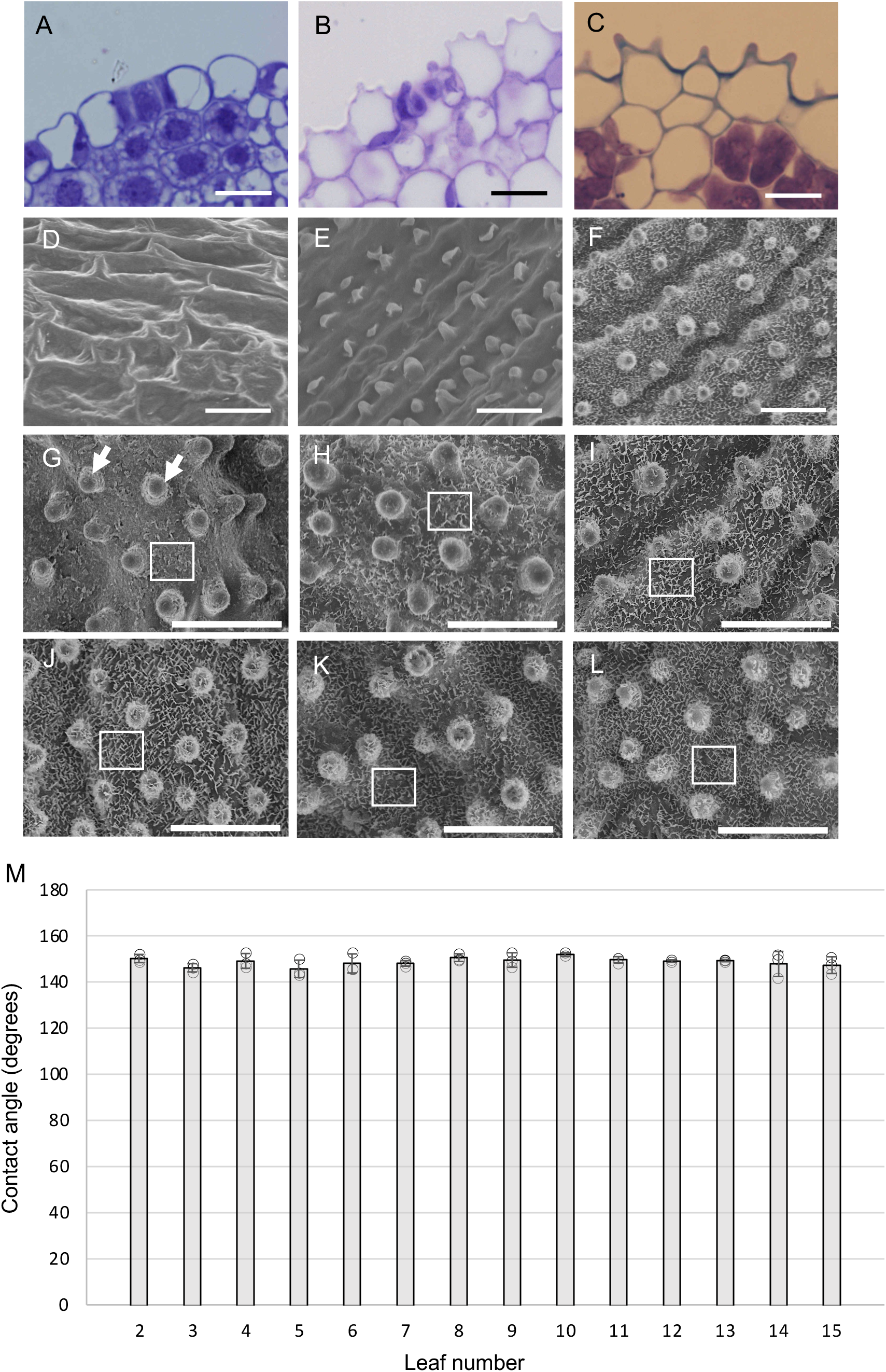
Changes in leaf surface structure during development and growth stages. **A–C**) Plastic sections from the basal to apical direction of the P4 leaf blade. **D–F)** Scanning electron microscopy images of the P4 leaf blade surface. The basal part (**A**, **D**); the central part (**B, E**); the apical part (**C**, **F**). **G–L**) Scanning electron microscopy images of the adaxial surface of the leaf blade. 2nd leaf (**G**); 3rd leaf (**H**); 4th leaf (**I**); 5th leaf (**J**); 6th leaf (**K**); 7th leaf (**L**). The white arrows in (**G**) indicate papillae. Scale bars in (**A–L**) = 10 µm. Note that the microstructure of epidermal wax crystals (white square in [**G–L**]) is altered in different leaves. (**M**) Contact angles of the adaxial leaf blades in various leaves. Error bars indicate standard deviation (*n* = 3).

A hallmark of phase change in maize development is that the amount of epidermal wax on the leaf surface decreases as the leaf position increases (Sylvester et al., 1990; Poethig, 1990). Wax accumulation may be important for water repellence and may also vary during the developmental stages or rice. Therefore, we measured contact angles on the adaxial surfaces of various leaf blades on the same plants. The contact angles were approximately 150° at all leaf positions and there were no significant differences (Fig. 2M). However, the microstructure of the wax crystals varied depending on the leaf position. Scaly wax was present on the 2nd and 3rd leaves, whereas reticulate wax was present on the 4th to 8th leaves (Fig. 3G–L). The shapes of the papillae on different leaves were not significantly different. Therefore, the microstructure of epicuticular wax varies with leaf position but this does not affect water repellence in rice.

**Figure 3.**
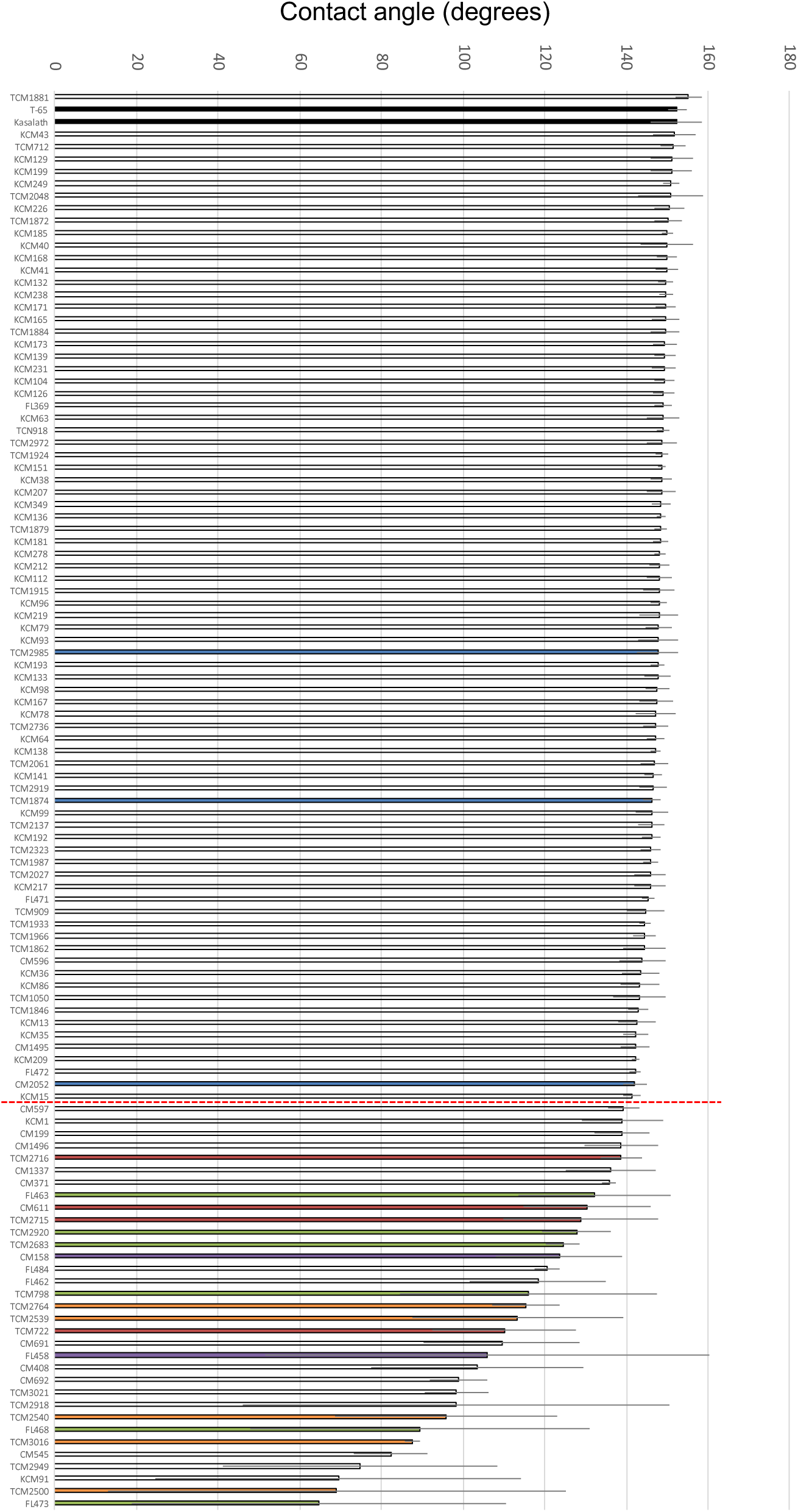
Contact angles of the adaxial leaf blades of the 4th leaf in potential mutant strains. Bars of the same color indicate mutant strains derived from mutations in the same gene. Black: wild-type; blue: *bgl*; red: *wsl4*; green: *drp7*; purple: *wsl2*; orange: *wsl3*. Error bars indicate standard deviation (*n* = 3). The red dotted line separates strains with contact angles of more than or less than 140°.

### Surface structure and water repellence of wetting-leaf mutants

Many rice mutant strains with a wetting-leaf phenotype have been collected by the National Bioresource Project (NBRP) (Oryzabase: https://shigen.nig.ac.jp/rice/oryzabase/). These probably include mutants with reduced water repellence. To characterize these potential mutants, we selected 113 strains that were described as having a wetting-leaf phenotype from the rice mutant stock maintained by the NBRP. We measured contact angles on the adaxial surface of the 4th leaf blade of 115 strains compared to wild-type (Japonica variety T-65 and the Indica variety Kasalath). When the mean values of the contact angles of these 115 strains were examined, all except one of the strains exhibited smaller contact angles than the two wild-type varieties. In addition, these strains could be separated into two groups. One group exhibited a small reduction in contact angle and little variation among individuals, whereas the other group exhibited a large reduction in contact angle and extensive variation among individuals (contact angle approximately 140°; Fig. 3).

We screened the potential wetting-leaf mutants using SEM and identified at least 20 strains with abnormal leaf surface microstructure (Fig. 4). These were separated into two types. The first type of mutant had no small papillae but normal epicuticular wax on the leaf surface (Fig. 4A–C). The second type had normal papillae but little or no epicuticular wax crystals (Fig. 4D–T). The first group of mutants included the three strains with contact angles > 140°. The second group of mutants all had contact angles of 140° or less, with the smallest contact angle being only 65° (Fig. 3). These results indicate that the rice wetting-leaf mutants had reduced water repellence but two different types of microstructure abnormality: no small papillae or insufficient accumulation of wax crystals. The reduced accumulation of wax crystals was more detrimental to water repellence than the absence of small papillae.

**Figure 4.**
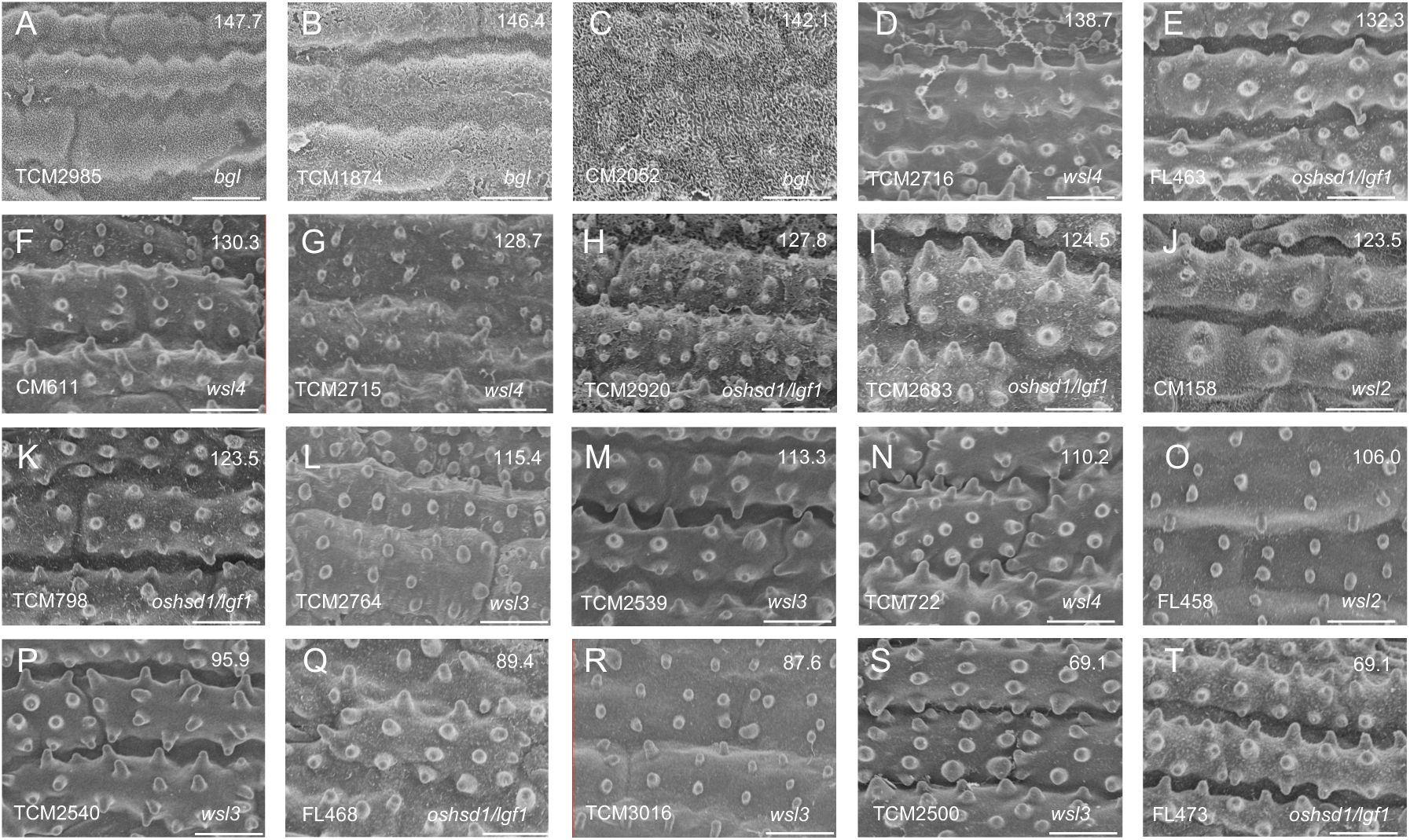
Scanning electron microscopy images showing the epidermal structure of 20 wetting-leaf mutants. **A–C**) The first type of epidermal structure: small papillae absent and normal wax crystals. **D–T**) The second type of epidermal structure: normal papillae but few wax crystals or none. For each panel, the contact angle (top right), strain name (bottom left), and mutant gene (bottom right) are indicated. Scale bars = 10 µm.

### Identification of the genes responsible for reduced water repellence in leaves

While investigating the genes responsible for reduced water repellence, we discovered that 1 of the 20 mutants, CM2052, had already been described as *bright green leaf* (*bgl*) mutant KL208, which exhibits altered leaf color and has no small epidermal papillae (Yoo et al., 2011). Therefore, we predicted that the mutations in TCM1874 and TCM2985 were also in the *BGL* gene, because these mutants also had no small epidermal papillae (Fig. 4A, B). We determined the DNA sequence of the *BGL* gene in both mutants and found that the same mutation generated a stop codon in the second exon in each case (Fig. 5A). This strongly suggests that TCM1874, CM2052, and TCM2985 all contained mutant alleles of the *BGL* gene.

**Figure 5.**
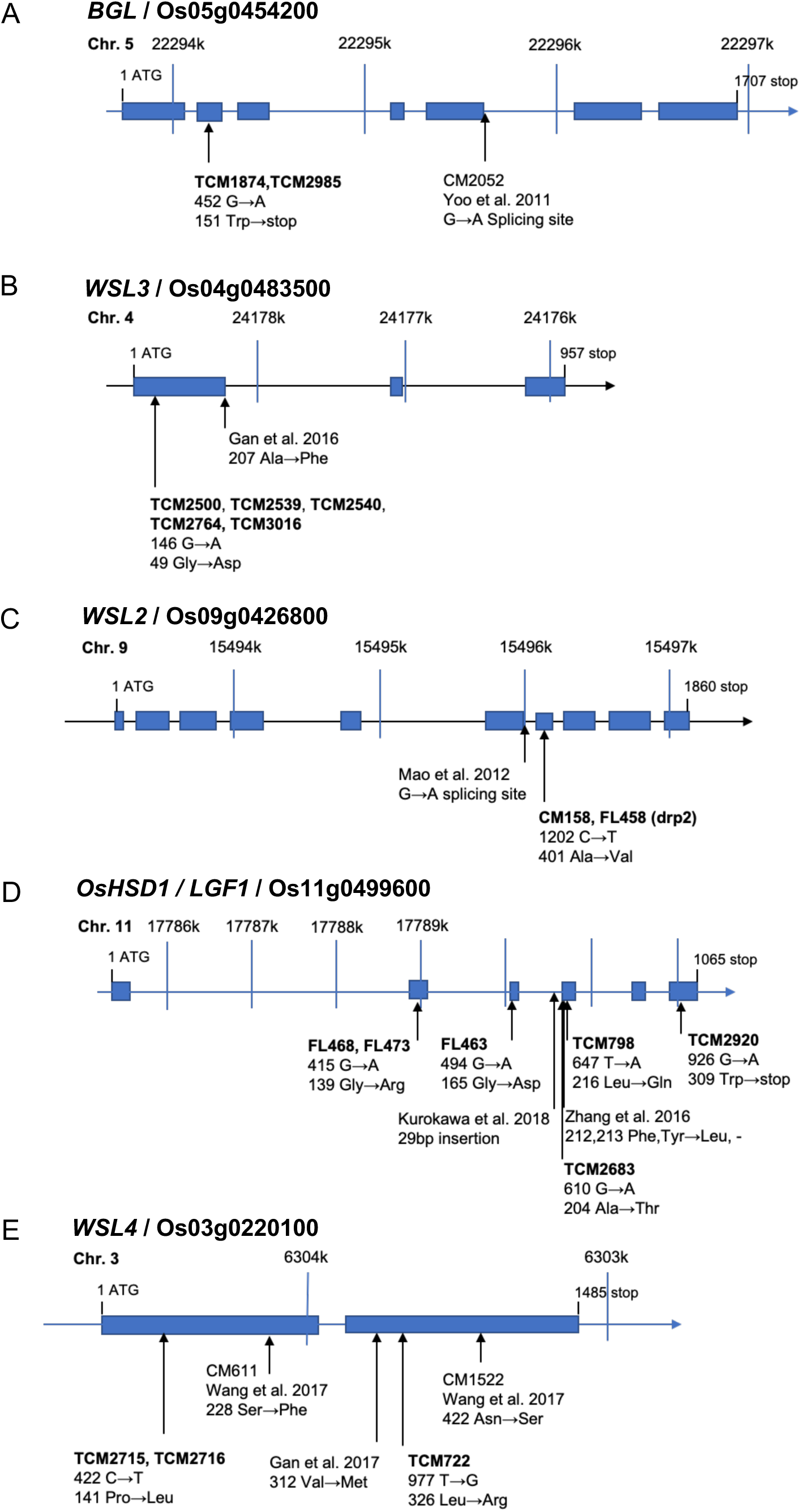
Genes and mutations involved in leaf water repellence. Mutant strains and mutations in *BGL*/Os05g0454200 (**A**), *WSL3*/Os04g0483500 (**B**), *WSL2*/Os09g0426800 (**C**), *OsHSD1/LGF1*/Os11g0499600 (**D**), and *WSL4*/Os03g0220100 (**E**). The mutant strains in bold are those identified in this study. The type and position of each mutation is shown, together with the resulting change in amino acid.

Of the remaining 17 strains, CM611 reportedly has a mutation in the *WSL3* gene, which encodes a KCR (Gan et al., 2016; Fig. 5B). Of the remaining 16 strains, 8 were crossed with the wild-type Indica variety Kasalath, and the F_2_ population was used to map the corresponding mutations. The mutations were mapped to the long arm of chromosome 4 (TCM2500, TCM2540, TCM2764, and TCM3016), the long arm of chromosome 9 (CM158 and FL458), the long arm of chromosome 11 (TCM2920), and the short arm of chromosome 3 (TCM2716). The following genes that are involved in wax biosynthesis are located in these four chromosomal regions: *WSL3* (Gan et al., 2016), *OsGL1–1*/*WSL2* (Qin et al., 2011; Mao et al., 2012), *OsHSD1/LGF1* (Zhang et al., 2016; Kurokawa et al., 2018), and *WSL4* (Gan et al., 2017; Wang et al., 2017). Therefore, we determined the DNA sequences of these genes in all 16 mutant strains and identified single nucleotide substitutions in one of the four genes in all strains (Fig. 5B–E). The reduced or absent accumulation of epidermal wax crystal phenotypes were very similar to those of mutants that had already been characterized (Qin et al., 2011; Mao et al., 2012; Gan et al., 2016; Zhang et al., 2016; Gan et al., 2017; Wang et al., 2017 Kurokawa et al., 2018), strongly suggesting that the 16 mutant strains with abnormal epicuticular wax were derived from mutations in four genes involved in wax biosynthesis.

The relationship between the mutations and the reductions in contact angle was analyzed. For mutant alleles of *BGL*, the contact angles of TCM1874, TCM2985, and CM2052 were 146.4°, 147.7°, and 142.0°, respectively (Fig. 3). Among the mutants of wax biosynthesis-related genes, mutations in *WSL3* that produced the same amino acid substitution were observed in TCM2500, TCM2539, TCM2540, TCM2764, and TCM3016 (Fig. 5B). However, the contact angles differed significantly from 69.1° in TCM2500 to 115.4° in TCM2764 (Fig. 3). Similar results were observed for the *WSL2* gene, in which the same mutation caused an amino acid substitution but the contact angle in CM158 was 123.5° but 106.0° in FL458 (Fig. 3 and 5C). Five different mutations in six mutant strains and two mutations in three mutant strains were found in *OsHSD1/LGF1* and *WSL4*, respectively, but all of these mutations produced amino acid substitutions (Fig. 5D, E). Overall, there appeared to be no association between the type of wax biosynthesis gene and the magnitude of the effect on contact angle: the magnitude of contact angle reduction could vary among mutant alleles of any of the four genes.

Identification of genes responsible for wetting-leaf phenotypes revealed that water repellence is regulated by at least one gene controlling papillae differentiation and four genes involved in the biosynthesis of epicuticular wax.

### Expression of wax biosynthesis-related genes

The expression patterns of the four wax biosynthesis genes (*WSL3*, *WSL2*, *OsHSD1/LGF1*, and *WSL4*) was investigated using the β-glucuronidase reporter gene, and staining corresponding to all four genes was observed in mature leaves (Qin et al., 2011; Mao et al., 2012; Gan et al., 2016; Zhang et al., 2016; Gan et al., 2017; Wang et al., 2017). However, epicuticular wax accumulation begins in leaf primordia at P4 (Fig. 2) and information regarding tissue specificity and the timing of expression is necessary to understand the regulation of epidermal wax accumulation. Contrary to our expectations, *in situ* hybridization using immature shoot tissue revealed expression of *WSL3* and *WSL4* (encoding a KCR and KCS, respectively) early in leaf development (Fig. 6A, B, G, H). *WSL3* and *WSL4* expression was detected not only in the L1 layer of P1 to P5 leaf primordia but also in the SAM. However, the expression pattern of *WSL2* differed from the other genes and signal was observed at P5 and P6, as well as in vascular bundles and sclerenchyma (Fig. 6C, D). The expression of *OsHSD1/LGF1* was observed in the L1 layer on the abaxial surface of the P5 leaf sheath (Fig. 6E, F).

**Figure 6.**
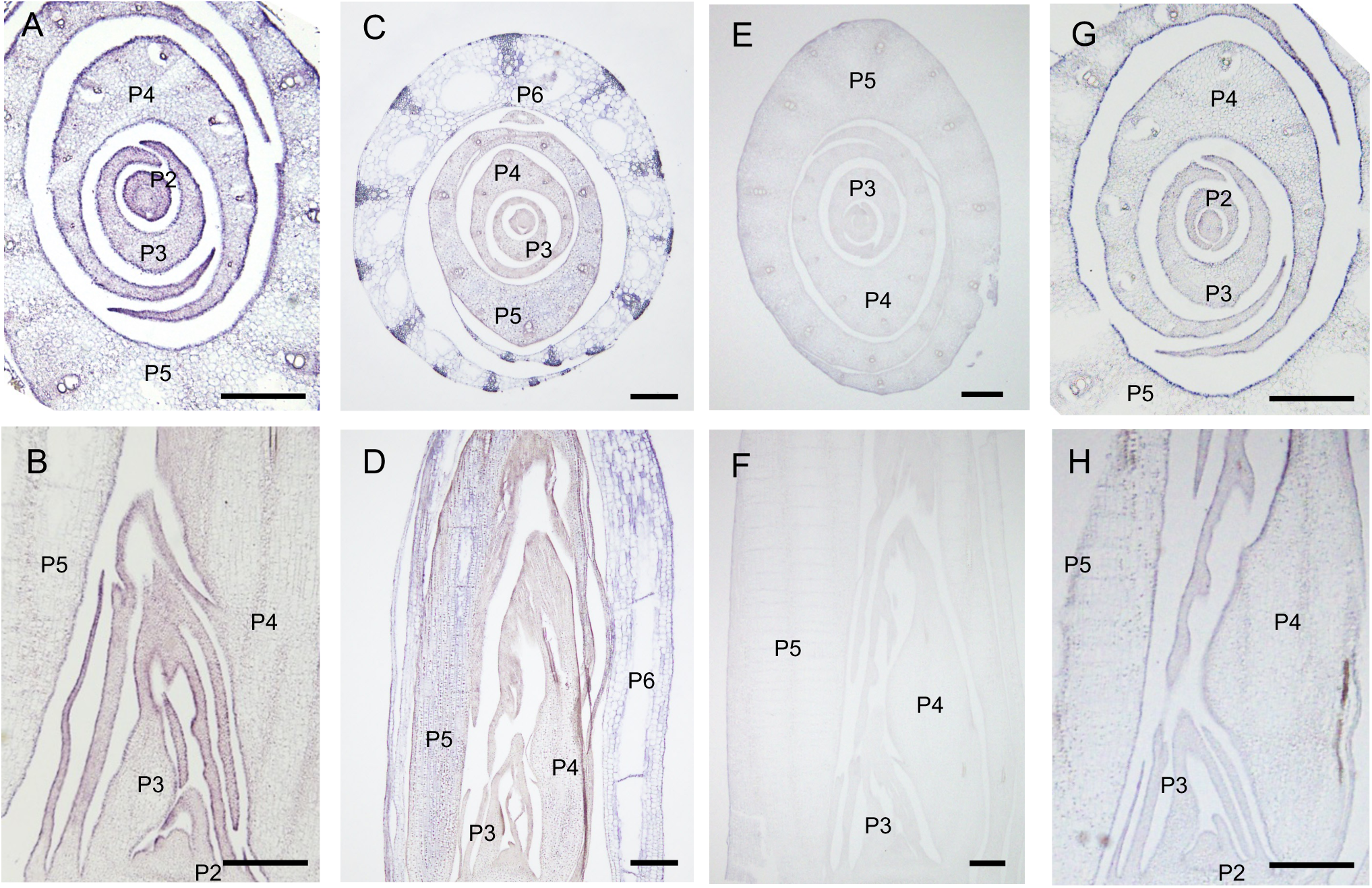
Spatial expression pattern of wax biosynthesis-related genes highlighted by *in situ* hybridization. Expression of *WSL3*/Os04g0483500 (**A** and **B**), *WSL2*/Os09g0426800 (**C** and **D**), *OsHSD1/LGF1*/Os11g0499600 (**E** and **F**), and *WSL4*/Os03g0220100 (**G** and **H**). Cross sections and longitudinal sections of wild-type seedlings 17 (**C** and **D**) and 25 (**A, B, E, F, G, H**) days after germination. The leaf stage (Px) is labeled. Scale bars = 200 µm.

Next, to determine whether the expression of these genes was affected by deficiencies in wax biosynthesis or papillae formation, expression levels were analyzed by qRT-PCR in each mutant (Supplementary Fig. S1). The expression of *WSL2* was reduced in the *wsl3* mutant, but there were no significant changes in the expression of any of the other genes in any of the mutants. These results suggest that the four genes involved in the biosynthesis of cuticular wax have different expression patterns, although the corresponding mutants show similar abnormalities in leaf wax accumulation. In addition, the expression level of each gene is largely unaffected by mutations in the other genes.

### Diversity of water repellence among rice varieties

To investigate diversity among rice cultivars and related species, we evaluated water repellence and epidermal structure in cultivated rice and wild *Oryza* species. First, we measured the contact angles of 55 accessions from the WRC, which includes cultivated rice and genetically diverse accessions maintained by NARO Genebank (https://www.gene.affrc.go.jp/databases-core_collections_wr_en.php; (Tanaka et al., 2020; Supplementary Table S1). More than half of the accessions had contact angles exceeding 150° and the smallest contact angle was 143° (WRC3), indicating that most of these cultivated rice accessions showed superhydrophobicity (Supplementary Fig. S2). Observations of leaf surface structure using low-vacuum SEM showed that all of the accessions were similar (Supplementary Fig. S3). This suggests that superhydrophobicity is conserved among rice cultivars and is an essential trait for rice cultivation around the world.

Next, to understand the origins of superhydrophobicity in cultivated rice, we investigated water repellence in wild *Oryza* accessions maintained by the National Institute of Genetics, Japan (https://shigen.nig.ac.jp/rice/oryzabase/locale/change?lang=en). To identify accessions with reduced water repellence we used a “shower test” screening procedure. Water was showered onto the leaves of approximately 400 wild *Oryza* accessions and we observed whether droplets of water remained on the leaf surfaces. A total of 31 “shower test positive” accessions were selected and we sampled mature leaf blades from these accessions to compare with WRC1 (Nipponbare) as a representative of cultivated rice (Supplementary Table S2). Unlike the WRC results, the contact angles of selected wild *Oryza* accessions suggested diverse degrees of water repellence: 25 of the 31 accessions had contact angles below 140°, which was significantly lower than the contact angles observed in cultivated species (Fig. 7). Observations of leaf surface structure using low-vacuum SEM revealed that several accessions with small contact angles had significantly lower papillae densities than the other accessions (Fig. 7C–H, Supplementary Fig. S4). Then, we investigated the relationship between the papillae density and the contact angles in the leaves of 32 accessions. The results showed a correlation, but some accessions had greater papillae densities than cultivated rice accessions with smaller contact angles (e.g., W1453; Fig. 7D). This suggests that factors other than papillae density may be responsible for reduced water repellence in wild *Oryza* species.

**Figure 7.**
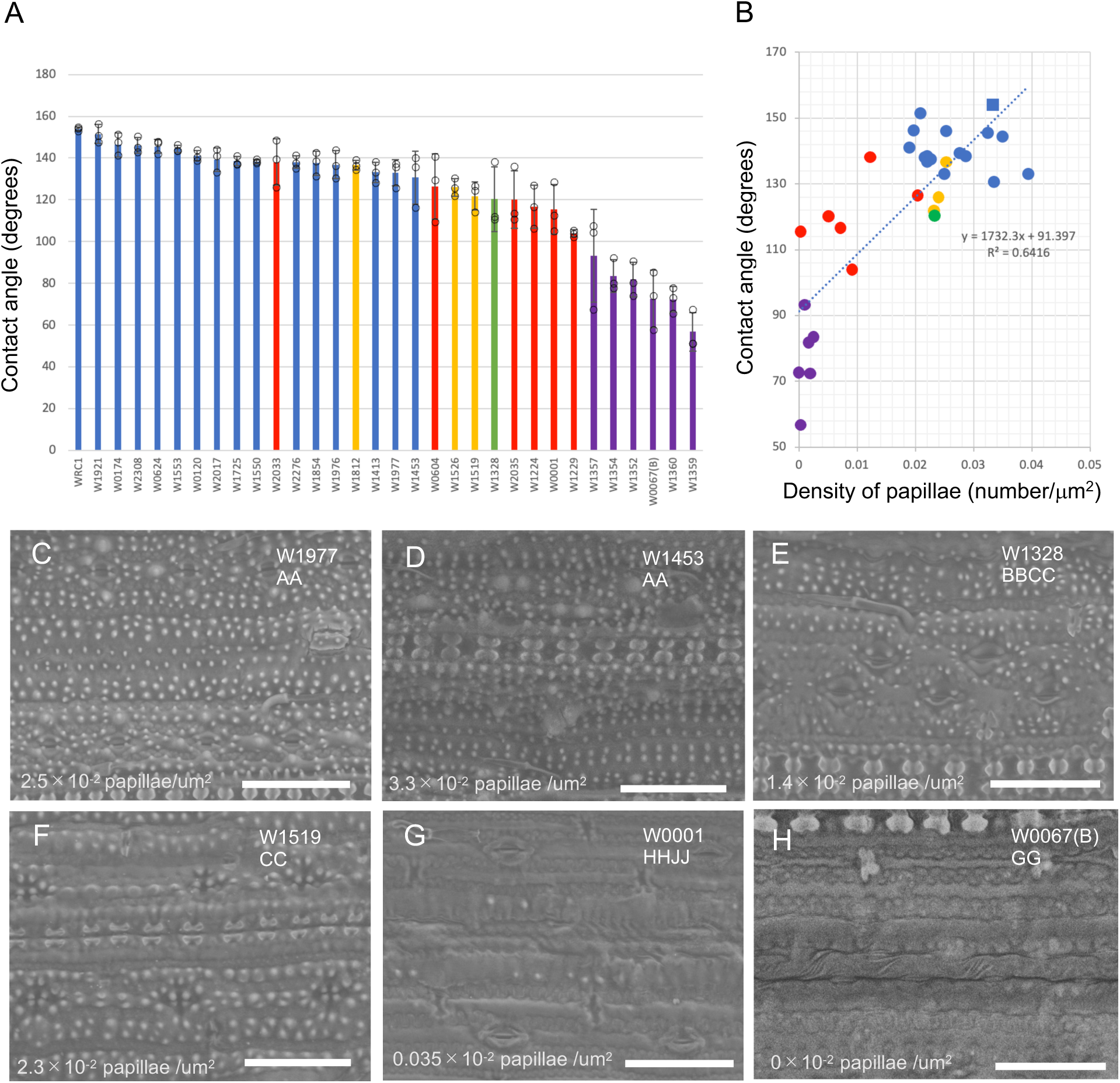
Water repellence, epidermal structure, and genome type in wild *Oryza* species. **A)** Contact angles of 31 wild *Oryza* accessions selected by shower tests and WRC1 (Nipponbare) as a representative of cultivated rice. Bars of the same color indicate the same genome type. Blue: AA; orange: CC; red: HHJJ; green: BBCC; purple: GG. Error bars indicate standard deviation (*n* = 3). **B)** Correlation between contact angle and papillae density. Plots of the same color indicate the same genome type, as described above. The square represents WRC1. **C–H)** The epidermal structure of several wild *Oryza* accessions revealed by low-vacuum scanning electron microscopy. For each panel, accession names and genome types are indicated in the top right corner and papillae density is shown in the bottom left corner. Scale bars = 50 µm.

The selected accessions of wild *Oryza* species could be classified into the following genome types: 15 AA (13 *O*. *rufipogon* and 2 *O*. *longistaminata*), 3 CC (2 *O*. *eichingeri* and 1 *O*. *rhizomatis*), 1 BBCC (*O*. *minuta*), 6 HHJJ (4 *O*. *ridleyi* and 2 *O*. *longiglumis*), and 6 GG (5 *O*. *meyeriana* and 1 *O*. *granulata*) species. The AA genome accessions had mean contact angles of approximately 140° and tended to have greater papillae densities than other genome types. On the other hand, HHJJ genome accessions had smaller contact angles and lower papillae densities, whereas the BBCC and CC accessions were phenotypically intermediate. Unlike other semi-aquatic species, the GG genome species that inhabit shady forests had extremely small contact angles and low papillae densities, and W0067(B) had almost no papillae. In addition, SEM showed no obvious wax crystals on the adaxial leaf surface of W0067(B) (Supplementary Fig S5). Therefore, most of the AA genome wild *Oryza* accessions that are most closely related to cultivated rice have high papillae densities and good water repellence. However, the *Oryza* genus includes some species with low papillae densities and limited water repellence, such as some of the GG genome accessions.

## Discussion

There are four different epidermal regions in rice leaves: the adaxial/abaxial leaf blade and the adaxial/abaxial leaf sheath (Itoh et al., 2005). Of these four regions, water repellence was greatest on the adaxial surface of the leaf blade and poorest on the adaxial surface of the leaf sheath. The poor water repellence of the latter compared to the other three regions is most likely due to differences in epidermal structure. That region is the only region that had neither papillae nor wax crystals. It is also the only region where the subsequent leaf remains in close contact, and it may not need a water-repellent structure because it interacts less with the external environment.

In maize, epidermal wax accumulates on the surface of lower leaves but is not present on higher leaves. This is considered one of the hallmarks of the juvenile-to-adult phase change in maize (Poethig, 1990). It is not clear whether the degree of water repellence differs between juvenile and adult maize leaves but this may be affected by differences in the chemical composition of the wax that occur during growth (Yang et al., 1993). On the other hand, there were no significant differences in contact angle in any rice leaves under our growth conditions (Fig. 2). This suggests that good water-repellent properties are maintained regardless of leaf position or growth stage. However, a comparison of epidermal structures revealed that the shape of the wax crystals covering the epidermis on the 2nd and 3rd leaves differs from that on mature leaves. There were scaly wax crystals on the 2nd and 3rd leaves but reticulate wax crystals on the mature leaves. Differences in wax composition may affect the shape of wax crystals (Pascal et al., 2019; Koch and Ensikat, 2008). Because the 1st- and 2nd-leaf stages are considered the juvenile phase in rice (Itoh et al., 2005), the composition of epicuticular wax may also differ between juvenile and adult leaves (Yang et al., 1993). This may affect the crystallization of epicuticular wax.

An analysis of water repellence and surface structure in wetting-leaf mutants revealed two types of mutant with different characteristics: one with normal epicuticular wax but lacking small papillae and one with normal small papillae but deficient in the accumulation of epicuticular wax. The first type of mutant was derived from mutations in the *BGL* gene, with *bgl* mutants previously characterized as luminous green plants that lack small papillae on their leaves (Yoo et al., 2011). We found that *BGL* is also involved in water repellence. The reduction in water repellence, which was associated with a reduction in contact angle of approximately 10° due to the loss of small papillae, was clearly smaller than that due to wax abnormalities. However, this change is significant for the physical properties of the leaf surface. A superhydrophobic surface is defined as one with a static contact angle greater than 150°, thereby creating a self-cleaning effect (Barthlott et al., 2017; Koch and Barthlott, 2009). If the surface of a rice leaf has normal epicuticular wax but lacks papillae, as in the *bgl* mutants, the contact angle cannot exceed 150°, resulting in the loss of superhydrophobicity. Therefore, papillae differentiation is necessary for rice plants to generate superhydrophobic leaves. Only a few grass species have papillae on their leaf surfaces and these are not necessarily closely related phylogenetically (Prasad et al., 2011).

Consequently, rice probably acquired or maintained papilla formation during the evolution of the *Poaceae*, fostering superhydrophobic characteristics suitable for semi-aquatic environments. How papillae differentiate remains unclear but papillae protrusions occur by the P4 stage of leaf development (Fig. 2) together with other changes in epidermal cells. Understanding the molecular mechanisms involved in this process may provide opportunities to manipulate water repellence by modifying surface structures.

The second type of mutant with reduced water repellence was derived from mutations in four wax synthesis genes. One characteristic of these mutants is large variation in contact angle among individual plants. The explanation for this remains unclear but a decrease in water repellence may reflect an increase in the contact area between a surface and a water droplet, which makes the contact angle difficult to measure. In addition to the variation in contact angle among individual plants, mean angles varied among the mutant strains. Perhaps the effects on water repellence vary depending on gene function and the strength of mutant alleles. In our analysis, however, most of the mutations in the four wax-synthesis genes produced changes in the amino acids, and there were no obvious null alleles. Therefore, we were unable to quantify the contribution of each gene to water repellence. In addition, even mutant strains with the same mutation showed variation in mean contact angle. Perhaps the differences between mutant strains are influenced by differences between individual plants and background mutations in the mutant strains. Notably, the mutant exhibiting the smallest contact angle among the 113 strains was FL473. This strain has a mutation in *OsHSD1/LGF1*, which is involved in C30 primary alcohol synthesis (Kurokawa et al., 2018). Interestingly, even in this mutant a few wax crystals were observed (Fig. 4T), whereas some mutants with greater water repellence had no visible wax crystals (Fig. 4M, N). Other researchers have noted the presence of some wax crystals in *OsHSD1/LGF1* gene mutants (Zhang et al., 2016; Kurokawa et al., 2018). Therefore, water repellence depends not only on the level of wax crystal accumulation on the epidermis but also on the composition of the wax film covering the cuticular surface.

We investigated the expression of four genes involved in epidermal wax synthesis and found that *WSL3* and *WSL4*, which are part of the FAE complex, are expressed in epidermal cells during early leaf development, including in the L1 layer of the SAM. This expression pattern was similar to that of *ONI1* and *ONI2*, which belong to the same KCS gene family as *WSL4* (Ito et al., 2011; Tsuda et al., 2013; Gan et al., 2017; Wang et al., 2017). Mutants of *ONI1* and *ONI2* were seedling lethal and showed developmental abnormalities including leaf fusion due to loss of function of the L1 layer (Ito et al., 2011; Tsuda et al., 2013). In contrast to *oni1* and *oni2*, all *wsl4* mutants were viable and no abnormalities were observed during early development. Although different KCS enzymes may have specific substrates (Batsale et al., 2021), *WSL4* may exhibit functional redundancy with *ONI1/ONI2* during early leaf development. Alternatively, it may have functions other than epidermal wax synthesis (Gan et al., 2017). Interestingly, two genes involved in modification after VLCFA biosynthesis showed expression patterns that differed from those of *WSL3* and *WSL4*. Expression of *OsHSD1/LGF1* was observed in the leaf epidermis during late development, whereas *WSL2* was expressed not only in the epidermis but also in the primordia of inner leaves (Qin et al., 2011). This suggests that *WSL2* functions not only in epicuticular wax synthesis but also in inner leaf tissues (Qin et al., 2011). In addition, mutations in each of four wax synthesis genes and in papilla-forming genes did not affect the expression of the wax synthesis genes. Therefore, under our experimental conditions, the expression of these four genes was regulated independently of changes in wax synthesis and papilla formation, although feedback regulation does reportedly occur among the wax synthesis genes (Mao et al., 2012).

No significant differences in contact angle or surface structure were observed among cultivated rice varieties from the WRC. This suggests that water repellence is conserved in cultivated rice. Among the WRC accessions, WRC51 is an upland rice cultivar but exhibits a similar degree of water repellence to that of other paddy rice accessions. Thus, water repellence may also be advantageous for varieties grown in dry conditions. On the other hand, leaf surface structure and the degree of water repellence did vary among wild *Oryza* accessions. Interestingly, AA genome accessions including *O*. *rufipogon*, thought to be ancestral to cultivated rice (*O*. *sativa*; Huang et al., 2012), tended to be more water repellent and had higher densities of small papillae. Because the population we used in our shower test screen contained more than 200 *O*. *rufipogon* accessions, many of the shower test negative accessions probably had very good water-repellent properties. Therefore, leaf superhydrophobicity was probably not selected for during the domestication of rice, but was already present in the ancestral species and inherited by *O*. *sativa*. The GG genome species are the only group adapted to forest rather than semi-aquatic environments (Shi et al., 2020). These GG genome species had very low papillae densities. Therefore, the differentiation of papillae might be adaptive for *Oryza* species in semi-aquatic environments. On the other hand, sometimes there was no relationship between papillae density and water repellence (e.g., WRC1 vs. W1453). In addition, significant differences in water repellence were found even between lines lacking papillae (e.g., *bgl* mutants vs. W0067[B]). W0067(B) did not exhibit the accumulation of wax crystals that was observed in cultivated rice (Supplementary Fig. S5). Therefore, differences in the accumulation and composition of wax may influence water repellence in various wild *Oryza* species.

Many studies have sought to predict what surface structures are important for water repellence, including superhydrophobicity (Neinhuis and Barthlott, 1997; Koch and Barthlott, 2009; Bhushan, 2012; Barthlott et al., 2017). However, there are few experimental reports describing how changes in surface structure affect water repellence in a single species or in closely related species. Our study reveals part of the structural, genetic, and adaptive basis for superhydrophobicity in rice leaves. Water repellence is the most fundamental physical property of plants, and manipulating this property may provide great opportunities for plant breeders (Jolliffe et al., 2023). For example, crops with superhydrophobicity may be more disease resistant and tolerant to drought or flooding. Conversely, reducing water repellence in leaves may facilitate the uptake of fertilizers and pesticides from the leaf surface (Taylor, 2011). Our results describing some of the key genes, gene expression patterns, and interactions between wax synthesis genes may be useful for genetic engineering of leaf repellence. Notably, the development of superhydrophobic materials that mimic the lotus leaf surface is one of the best-known examples of biomimetics (Koch and Barthlott, 2009; Bhushan, 2012; Barthlott et al., 2017), and a deeper understanding of plant superhydrophobicity could be applied to facilitate the development of high-performance products.

## Materials and Methods

### Plant materials

Taichung 65 (T-65) and Kasalath were used as representative varieties of rice (*Oryza sativa* L.). Wetting-leaf mutants derived from N-methyl-N-nitrosourea-treated accessions stored at Kyushu University were identified by text screening using Oryzabase (http://www.shigen.nig.ac.jp/rice/oryzabase/). There was a total of 113 strains, including mutant strains from the Kinmaze, T-65, and Kitaake background lines (CM, TCM, and KCM) and marker gene accumulation lines (FL).

For diversity analysis of rice cultivars, 55 of the 69 accessions from the World Rice Core Collection (WRC) stored at the Genebank for Agricultural Biological Resources of the National Agriculture and Food Research Organization (NARO) were used (Supplementary Table S1). The WRC collection covers 90% of allelic diversity (Kojima et al., 2005; Tanaka et al., 2020; https://www.gene.affrc.go.jp/databases-core_collections_wr.php).

For diversity analysis of wild *Oryza* species, we used 31 accessions from the wild *Oryza* species collection (https://shigen.nig.ac.jp/rice/oryzabase/strain/wildCore/list) stored and maintained at the National Institute of Genetics in Japan (Supplementary Table S2).

### Growth conditions

The wild-type and mutant plants used to measure contact angles and analyze surface structure were grown under natural conditions. For *in situ* hybridization and quantitative real-time polymerase chain reaction (qRT-PCR), wild-type and mutant seeds were sterilized and sown on soil or Petri dishes and subsequently grown in a growth chamber with a 16 h light (30°C)/8 h dark (25°C) photoperiod.

Wild *Oryza* species grown under natural conditions in a greenhouse or a field at the National Institute of Genetics in Mishima were screened using a “shower test” in early July. Leaf blades that had developed and emerged completely from the leaf sheath but were not senescent were sampled.

### Measurement of contact angles

The center of the leaf blade or leaf sheath was cut to an appropriate size, fixed on a glass slide using double-sided tape with the abaxial surface facing upward, and placed on the stage of a contact angle meter (LSE-ME3 Nick, Saitama, Japan). A syringe was used to apply 1 µL water to the leaf surface and the contact angle was measured for 5 s. Leaf tissue was collected from the same position in three separate plants. The adaxial surface of the center of the leaf blade was measured three times per leaf.

### Observation of surface structure

Sampled leaf blades were immersed in 4% paraformaldehyde solution followed by overnight fixation. After a graded series of tert-butyl alcohol washes, samples were immersed in 100% tert-butyl alcohol and frozen at –20°C. The frozen samples were freeze-dried using a lyophilizer (JFD-320; JEOL, Ltd, Tokyo, Japan). Samples were prepared for scanning electron microscopy (SEM) under a stereomicroscope and secured to the stage using carbon tape. A platinum ion coating was applied for 90 s using an E-1030 ion sputter coater (E-1030; Hitachi, Tokyo, Japan). Leaf surface structure was observed via SEM (S-4800; Hitachi). To view the surface structure of WRC lines and wild *Oryza* species, we applied low-vacuum tabletop SEM using the Miniscope TM1000 (Hitachi) and JCM-7000 NeoScope instruments (JEOL). Sampled leaf blades were glued to the stage using carbon tape under a stereomicroscope, and leaf surface structure was observed without further treatment.

### Preparation of plastic sections

Leaf samples were cut under a stereomicroscope, immersed in 4% (w/v) paraformaldehyde and 1% Triton X-100 in 0.1 M sodium phosphate buffer, and fixed overnight. The fixed samples were sequentially washed with ethanol (30%, 50%, 70%, 90%, and 100%) and then immersed in Technovit 7100:100% ethanol (1:1) and 100% Technovit 7100 (Heraeus Kulzer, Hanau, Germany). Samples were hardened by adding Technovit 7100:Hardener II (15:1). Embedded samples were cut into sections 3 µm thick using a microtome. Sectioned samples were stained with 0.05% toluidine blue and observed under an optical microscope.

### Identification of genes responsible for the wetting-leaf phenotype

To map TCM2500, TCM2540, TCM2764, TCM3016, CM158, FL458, TCM2920, and TCM2716, individual plants exhibiting the wetting-leaf phenotype were selected from an F_2_ population after crossing each mutant with Kasalath. DNA from selected individuals was analyzed for linkage using simple sequence repeat and cleaved amplified polymorphic sequence mapping markers (Supplementary Table S3). The chromosomal regions identified by these markers were analyzed for genes involved in wax biosynthesis or creating epidermal structure to identify candidate genes. DNA sequences and mutations for five candidate genes from the mutant strains were determined by Sanger sequencing.

### *In situ* hybridization

*In situ* hybridization was performed using antisense RNA probes. To synthesize the probes, cDNA fragments from three genes (*WSL2*, *OsHSD1/LGF1*, and *WSL4*) were amplified and cloned using the Zero Blunt TOPO PCR Cloning kit (Invitrogen, Carlsbad, CA, USA; Table S4). *WSL3* cDNA was obtained from a plasmid containing full-length cDNA (AK060512) provided by the National Institute of Agrobiological Sciences, Japan. These cDNA plasmids were used as templates and antisense RNA probes were synthesized using the DIG RNA Labeling kit (Roche Diagnostics GmbH, Mannheim, Germany).

Samples of T-65 plants were fixed using 4% (w/v) paraformaldehyde and 1% Triton X-100 in 0.1 M sodium phosphate buffer for 48 h at 4°C. Then they were dehydrated using a graded series of ethanol washes, immersed in 1-butanol, and embedded in Paraplast Plus (McCormick Scientific, Berkeley, MO, USA). The samples were cut into sections 8 µm thick using a rotary microtome and longitudinal and cross-sections containing shoot apical meristem (SAM) tissue were prepared. Hybridization and chromogenic reactions were performed as described by (Miya et al., 2025). After staining, the sections were mounted using Poly Mount (Polysciences, Inc., Warrington, PA, USA) and observed under a light microscope.

### qRT-PCR

We analyzed the expression levels of the *WSL3*, *WSL2*, *OsHSD1/LGF1*, and *WSL4* genes. Wild-type (T-65 and Kinmaze) and mutant (CM661 for *wsl4*, TCM2500 for *wsl3*, CM158 for *wsl2*, TCM798 for *oshsd/lgf1*, and TCM2985 for *bgl*) seeds were sterilized and sown on wet filter papers in Petri dishes. Whole shoots from two seedlings at 2 days after sowing were collected for each sample. Samples were frozen and RNA was extracted using TRIzol reagent (Invitrogen). The extracted RNA was treated with Recombinant DNase I (TaKaRa Bio, Inc., Shiga, Japan) and cDNA was synthesized using the High-Capacity cDNA Reverse Transcription kit (Life Technologies, Carlsbad, CA, USA). QRT-PCR was performed using the StepOne Real-Time PCR system (Life Technologies) with TaqMan Fast Universal PCR Master Mix and FAM-labeled TaqMan probes for each gene. The *OsRAD6* gene was used as an internal standard. The TaqMan probes and primers used for each gene are listed in Table S5. For all experiments, we analyzed three technical and four biological replicates.

### Measurement of papillae density

Images of leaf surfaces obtained using the tabletop Miniscope TM1000 (Hitachi) and JCM-7000 NeoScope SEM instruments were imported into ImageJ software (National Institutes of Health, Bethesda, MD, USA) to measure papillae densities. Densities were measured from three separate samples as the number of papillae in a predefined area.

## Acknowledgments

The mutant strains and wild *Oryza* accessions used in this study were obtained from the National Institute of Genetics, which is supported by the National Bioresource Project, MEXT, Japan. The World Rice Core Collection is distributed by NARO Genebank, Japan.

## Author contributions

J-i.I., T.Y. and Y.S. designed the research; A.H., S.A., Z.T., M.M., T.Y. and J-i.I. performed the experiments; A.H., S.A. and J-i.I. analysed the data; and J-i.I. wrote the manuscript. All authors offered suggestions on various drafts of the manuscript.

## Supplementary data

**Supplementary Figure S1.** Expression of wax synthesis genes in wetting leaf mutants. Relative expression level of *OsHSD1/LGF1*, *WSL4, WSL2, WSL3*. A ubiquitin-conjugating enzyme gene, *OsRAD6,* was used as an internal control. Single asterisks indicate a statistically significant difference compared to WT (t-test, P < 0.05).

**Supplementary Figure S2.** Contact angles of the adaxial leaf blade of 4th leaf in 55 WRC accessions. Error bars indicate standard deviation (*n* = 3).

**Supplementary Figure S3.** Epidermal structure of WRC accessions by low-vacuum SEM. For each panel, the accession name is indicated in the top left. Scale bars = 50µm.

**Supplementary Figure S4.** Epidermal structure of wild *Oryza* accessions by low-vacuum SEM. For each panel, accession names and genome types are indicated in the top left corner and papillae density (papillae/µm^2^) is shown in the bottom left corner. Scale bars = 50 µm. Scale bars = 50 µm.

**Supplementary Figure S5.** Scanning electron microscopy images showing the epidermal structure of W0067(B) accession. **A**) Scanning electron microscopy image of the adaxial surface of the leaf blade of W0067(B) accession. **B**) an enlarged view of the surface. Scale bars = 50µm in (**A**) and 10µm in (**B**).

**Supplementary Table S1.** List of WRC (world rice core collection) accessions used in this study.

**Supplementary Table S2.** List of wild *Oryza* accessions used in this study.

**Supplementary Table S3.** List of primers for linkage analysis.

**Supplementary Table S4.** List of primers for making in situ hybridization probes.

**Supplementary Table S5.** List of primers for quantitative RT-PCR.

